# The silicon transporters (SITs) gene was found in *Synechococcus* sp. XM24

**DOI:** 10.1101/2024.06.21.599990

**Authors:** Yabo Han, Jun Sun

## Abstract

Silicate serves as a crucial nutrient for diatoms, which are capable of absorbing dissolved silicon from their surroundings through silicon transporters (SITs), thus playing a significant role in the global ocean’s silicon cycle. Recent studies have indicated that marine single-celled *Synechococcus* also has the ability to accumulate silicon. Given that the evolution of *Synechococcus* predates that of diatoms, it is postulated that *Synechococcus* may utilize transporters such as SITs found in diatoms, to absorb dissolved silicon in the ocean. This research delves into the silicon accumulation in *Synechococcus* sp. XM24, particularly focusing on two key aspects. Firstly, the study investigates the potential presence of SITs in *Synechococcus* sp. XM24 under the condition of Depleted-Repleted silicate. Subsequently, two gene sequences suspected to encode *SITs*, were identified, and their protein sequences and functions were successfully predicted, shedding light on their involvement in membrane transport processes.

## Introduction

Silicate is a crucial nutrient for diatoms, as its absence halts cell division and leads to their demise. Diatoms absorb silicon in the form of orthosilicic acid (Si(OH)_4_) (Amo and Brzezinski, 1999) and transport it cooperatively with Na^+^, with a ratio of Si(OH)_4_ to Na^+^ concentration at 1:1 (Bhattacharyya and Volcani, 1980). Specific silicon transporters (SITs) facilitate the absorption of orthosilicic acid into the cell (Bäuerlein, 2000), initially discovered in diatoms (Hildebrand et al., 1997). Numerous SITs gene sequences, exceeding 400, have been identified across 36 genera based on diatom studies (LUPAS et al., 1991; Hildebrand et al., 1998; Armbrust et al., 2004; Thamatrakoln et al., 2006; Alverson, 2007; Dilkes et al., 2009; Curnow et al., 2012; Kang et al., 2015; Marchenkov et al., 2016; Marchenkov et al., 2018). Silicic acid serves as a significant pH buffer in diatom cells, crucial for maintaining stable acid-base balance. Its presence promotes the conversion of bicarbonate, providing diatoms with increased carbon dioxide resources to enhance growth and metabolic activities (Mann, 1999; Milligan and Morel, 2002). This mechanism enhances carbon dioxide absorption efficiency, improves photosynthesis, and accelerates biomass synthesis and energy capture in diatoms.

Cyanobacteria, a prevalent microorganism in various natural environments, is found in diverse water bodies, soils, and organisms. With a fossil record dating back approximately 2.7 billion years, cyanobacteria have played a significant role in Earth’s evolutionary history (BUICK, 1992; Brasier et al., 2002). Among marine cyanobacteria, *Synechococcus* stands out as a prominent group, first identified in 1979 (WATERBURY et al., 1979)_◯_ Characterized as small, single-celled algae ranging from 0.5 to 2 microns in diameter with a spherical shape (Johnson and Sieburth, 1979; Binder et al., 1996). *Synechococcus* is notable in marine ecosystems. Traditionally, diatoms have been credited with driving the global ocean’s silicon cycle due to their role in transporting dissolved silicon from the surface to the deep ocean. This understanding has long underpinned research on silicon cycling (Struyf et al., 2009), a key aspect of studying the silicon-carbon coupling cycle (Jun and Yuqiu, 2018). Diatoms, widely distributed in the ocean, are recognized for their siliceous shells that contribute significantly to the silicon cycle (Jun et al., 2016). However, further investigations into marine ecosystems have revealed challenges in solely attributing silicon and carbon production imbalances in oligotrophic oceans to diatoms. Recent studies have highlighted the dominance of picophytoplankton such as *Synechococcus* in primary productivity within oligotrophic regions, significantly influencing the export of carbon particles to the deep ocean (Richardson and Jackson, 2007; Lomas and Moran, 2011; Wei et al., 2022). This discovery poses a significant challenge to the established role of diatoms in the global marine silicon cycle (Jun and Yuqiu, 2018).

During the Precambrian era, phytoplankton had low evolutionary density, with marine silicon concentration remaining around 1.4mM (Conley et al., 2017) and amorphous silicon saturation at 2mM (Krauskopf, 1956; Iler, 1979). The discovery in 2012 that *Synechococcus* can accumulate silicon sparked significant interest among researchers, leading to a series of studies on silicon accumulation mechanisms (Baines et al., 2012; Tang et al., 2014; Deng et al., 2015; Guidi et al., 2016; Ohnemus et al., 2016; Brzezinski et al., 2017; Krause et al., 2017). Cyanobacteria do not exhibit a specific demand for silicon within the 1-120μM silicate concentration range. However, at 80-100μM silicate concentration, cyanobacteria demonstrate significant silicon enrichment akin to diatoms (Brzezinski et al., 2017). These findings suggest that during the Precambrian era with a 1.4mM silicate concentration, cyanobacteria may have developed a silicon accumulation mechanism before diatoms, potentially absorbing dissolved silicon in the ocean through transporters like SITs.

The current understanding of the mechanisms and environmental factors influencing silicon accumulation in *Synechococcus* remains incomplete. To investigate the presence of genes associated with silicon accumulation in *Synechococcus* and the impact of environmental variations on Biogenic Silica (BSi) accumulation, we conducted laboratory culture experiments and macrotranscriptomic analysis. Specifically, we established indoor culture experiments under Depleted-Repleted silicate conditions to elucidate the genetic regulation of silicate utilization in *Synechococcus*. Our study involved analyzing changes in silicon accumulation in *Synechococcus* under diverse environmental conditions through the assessment of physiological and biochemical parameters (Total BSi and New BSi) alongside macrotranscriptome sequencing. New BSi refers to the Si(OH)_4_ accumulated by *Synechococcus* and labeled by PDMPO.

## Materials and methods

### 1 The origin of *Synechococcus*

*Synechococcus* sp. XM24 originates from an algae species that was isolated by Professor Zheng from the coastal region of Xiamen (Zheng et al., 2018).

### 2 Culture method and condition

The medium was artificial seawater (Table S1). The light conditions were 100 μmol photons·m^-2^·s^-1^ and the culture volume was 400ml. Semi-continuous culture technique was employed, and the final concentration of PDMPO was 0.125μM (Leblanc and Hutchins, 2005). Three parallel samples were prepared for each condition for subsequent macrotranscriptome sequencing analysis. In the Depleted silicate group, all medium conditions were identical to the Repleted silicate group except for the absence of silicate, with a constant temperature of 25□.

### 3 Method for determination of BSi

Samples were collected once cell growth reached a stable state. Total BSi and New BSi simples were obtained by filtering 5ml of culture solution through 0.6μm PC membrane, then storing the membranes at -20□ in the dark. Sampling was conducted over 5 consecutive days to minimize errors. Total BSi was extracted using the hot alkali method (Azam et al., 1974), and its absorbance was measured at 812nm using silicon molybdenum blue spectrophotometry with an ultraviolet spectrophotometer. New BSi was extracted by combining silicon with PDMPO using the hot alkali method, followed by excitation it at 375nm with a fluorescence spectrophotometer to measure the emission spectrum of PDMPO at 530nm.

### 4 Macrotranscriptome sequencing

*Synechococcus* cells were filtered and collected with 0.2μm PC membrane, immediately frozen in liquid nitrogen, transferred to a -80□ refrigerator for storage, and sequenced as soon as possible. The macrotranscriptome sequencing in this study was done by the company WEKEMO.

### 5 Gene sequence analysis, protein structure and function prediction

The differential gene sequence obtained by sequencing was extracted and named, and a differential gene bank was established for comparison with the reference genome to screen out the differential genes that could be successfully matched with the reference genome, and these differential genes were considered to belong to *Synechococcus* sp. XM24. TBtools (Chen et al., 2023) was used for sequence screening, and the obtained sequences were translated into protein amino acid sequences using Seqkit (Zou et al., 2016). The protein structure and function were predicted using the HelixFold model (Wang et al., 2022) of PaddleHelix platform.

### 6 AutoDock4 molecular docking simulation

The small molecule used in this study is Orthosilicic Acid, which contains Si atoms. Si atoms are not commonly used in molecular docking, so the parameters of Si atoms are not included in the parameter file of AutoDock4 (Morris et al., 2009). Locate the AD4_parameters.dat file and add Si element parameters to it. Before running the software, add parameter_file AD4_parameters.dat to the first line of the GPF and DPF files respectively to enable AutoDock4 to run normally.

## Results and Discussion

### 1 Change of silicon accumulation of *Synechococcus* sp. XM24 under Depleted-Repleted silicate conditions

The cells were cultured individually in Depleted and Repleted silicate media under identical light conditions, maintaining consistent cell states through semi-continuous culture. BSi samples were collected over 5 consecutive days post stabilization of cell status, and data were analyzed using the T-test method. The results, depicted in Figure 1, show concentrations of Total BSi and New BSi in (A) and (B), respectively. In (A), the Repleted silicate group exhibited significantly higher levels compared to the Depleted silicate group (*, P < 0.01), indicating that cells could adapt to both living conditions at the same time, and the silicate content in cells would change with the change of silicate content in the environment. Notably, Depleted silicate did not entail complete removal of silicate from the culture medium. Silicate is inevitably brought into the preparation of various experimental reagents and culture containers. Therefore, “Depleted silicate” here means that no additional silicate is introduced, and the background silicate content is the same (Werner, 1977). In (B), New BSi levels were significantly elevated in the Depleted silicate state compared to the Repleted silicate state (*, P < 0.01), indicating an influence of external silicate levels on cellular silicate utilization. The increase in New BSi in the Depleted silicate group may be attributed to physiological requirements, such as new silicon-containing cell wall formation, potentially accelerated under environmental stress. This acceleration could prompt heightened silicate absorption from the extracellular environment, augmenting New BSi levels.

**Figure1.**
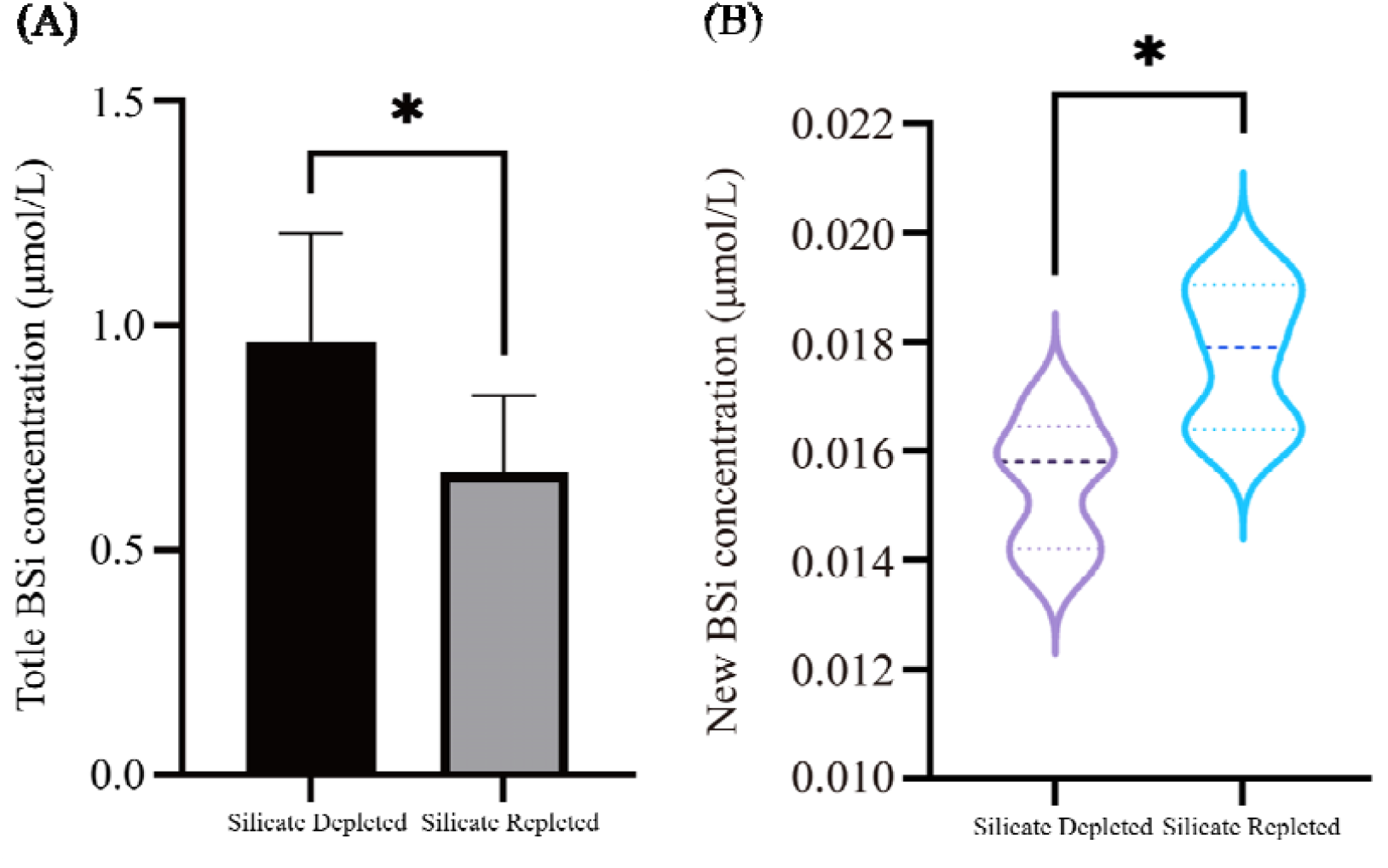
Under Depleted-Repleted conditions, the amount of silicon accumulated in the cell changes. (A) represents the concentration of Total BSi; (B) represents the concentration of New BSi. *, P < 0.01, indicating significant difference.

### 2 Macrotranscriptomic analysis of *Synechococcus* sp. XM24 under Depleted-Repleted silicate conditions

The TBtools software was utilized for sequence comparison in the process of sequence screening. The Blast Zone module was employed to create local databases by importing reference genome sequences and differential gene sequences. Subsequently, a comparison was made between these databases to identify the differential gene sequences of *Synechococcus* sp. XM24, resulting in the identification of 60 differential gene sequences. These sequences were then translated into protein amino acid sequences using Seqkit software. Based on their lengths, the sequences were categorized into four groups: less than 100, 100-1000, 1000-2000 and greater than 2000, comprising 6, 26, 13 and 15 sequences respectively. Following a reference to the protein sequence length of SITs in diatoms, analysis was focused solely on the 26 protein sequences falling within the 100-1000 length range. The HelixFold protein prediction model was employed to predict the structures of these 26 proteins, and only proteins with pLDDT (per-residue local distance difference test) scores exceeding 70 were retained. The pLDDT score, ranging from 0 to 100, serves as a confidence measure estimated by the model, with higher scores indicating a greater likelihood of the model closely resembling the true protein structure (Very high confidence: over 90; High: 70-90; Low: 50-70; Very low: below 50). Subsequently, 3 proteins with scores surpassing 70 were identified. Finally, the protein function prediction module within the PaddleHelix platform was utilized to predict the function of each protein. Two proteins A07_TRINITY_DN753_c0_g1 and e_TRINITY_DN668_c0_g1 successfully revealed their functions, whereas e_TRINITY_DN14139_c0_g1 did not. The gene and protein sequences for these proteins can be found in Supplementary Material 2. Figure S1-S2 display the locations of differentially expressed genes in relation to the reference genomes. Tables 1 and 2 present the Gene Ontology (GO) prediction results.

**Table1.**
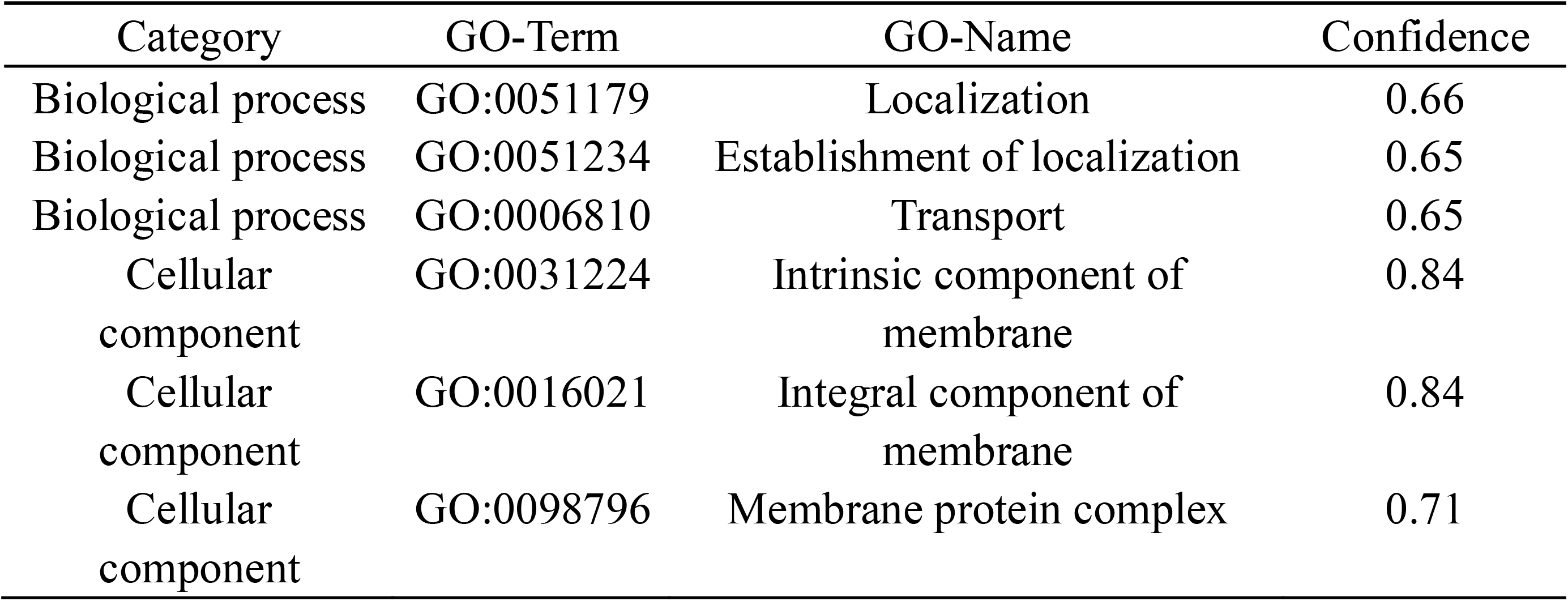
A07_TRINITY_DN753_c0_g1 protein function prediction results.

| Category | GO-Term | GO-Name | Confidence |
| --- | --- | --- | --- |
| Biological process | GO:0051179 | Localization | 0.66 |
| Biological process | GO:0051234 | Establishment of localization | 0.65 |
| Biological process | GO:0006810 | Transport | 0.65 |
| Cellular component | GO:0031224 | Intrinsic component of membrane | 0.84 |
| Cellular component | GO:0016021 | Integral component of membrane | 0.84 |
| Cellular component | GO:0098796 | Membrane protein complex | 0.71 |

**Table 2.**
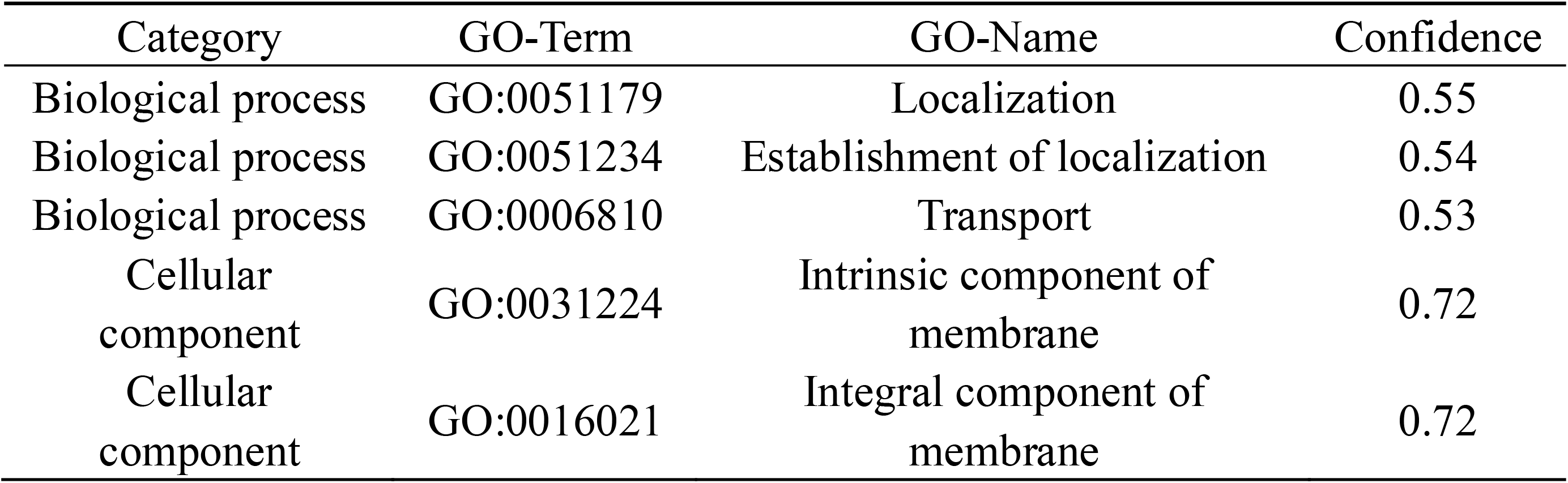
e_TRINITY_DN668_c0_g1 protein function prediction results.

Figure S1-S2 displays the positions of two differential gene sequences within the reference genome, with corresponding base numbers and “-” denoting the complementary strand. The analysis reveals that these sequences are reverse complementary to the reference genome. Tables 1 and 2 present the predicted functional outcomes of proteins translated from these sequences. Prediction confidence is indicated, with a threshold of 0.5 for credibility. The proteins derived from both sequences are likely associated with cell transport in Biological processes and cell membrane composition in Cellular components. These findings suggest a probable involvement of these proteins in cellular silicate transport, specifically implicating A07_TRINITY_DN753_c0_g1 and e_TRINITY_DN668_c0_g1 in regulating this process. It is plausible that these proteins correspond to the *SIT* gene in *Synechococcus* sp. XM24.

### 3 Molecular docking

Proteins and small molecules were subjected to molecular docking simulation using AutoDock4. Two proteins identified earlier were employed as receptors, while Orthosilicic Acid (provided in Supplementary Material 2) served as the ligand for docking. The most stable structure obtained from the docking simulations is illustrated in Figures 2 and 3.

**Figure 2.**
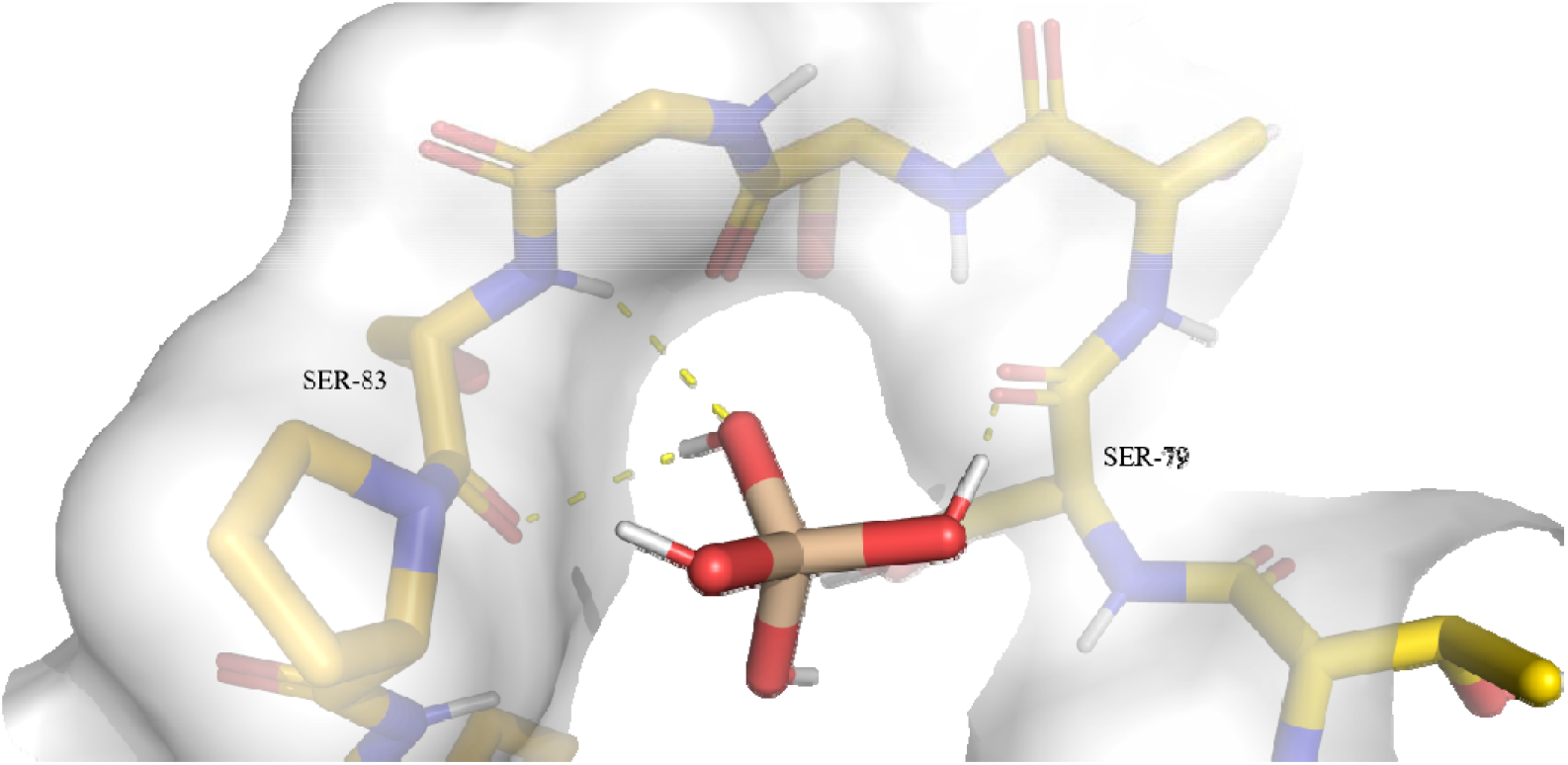
The docking result of A07_TRINITY_DN753_c0_g1 protein and Orthosilicic Acid. White for hydrogen atoms, red for oxygen atoms, blue for nitrogen atoms, yellow for carbon atoms.

**Figure 3.**
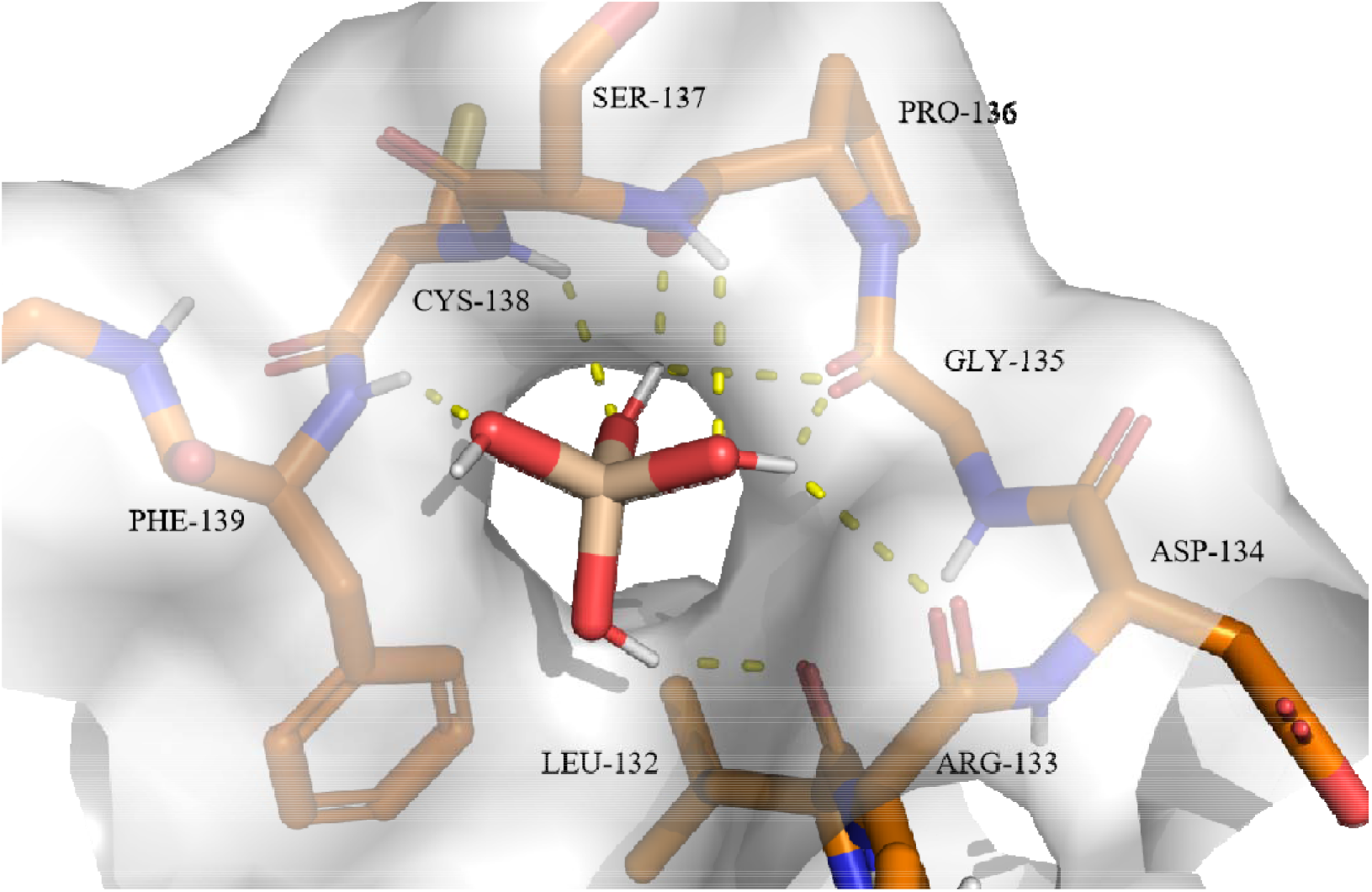
The docking result of e_TRINITY_DN668_c0_g1 protein and Orthosilicic Acid. White for hydrogen atoms, red for oxygen atoms, blue for nitrogen atoms, brown for carbon atoms.

The binding affinity between the A07_TRINITY_DN753_c0_g1 protein and Orthosilicic Acid is -2.18 kcal/mol, with the primary docking site identified at the hydroxyl group of Orthosilicic Acid and the protein residues SER-80 and SER-83, resulting in the formation of three hydrogen bonds. On the other hand, the binding affinity between the e_TRINITY_DN668_c0_g1 protein and Orthosilicic Acid is -3.31 kcal/mol, with the main docking site located at the hydroxyl group of Orthosilicic Acid and the protein residues LEU132, ARG-133, GLY-135, PRO-136, SER-137, CYS-138 and PH-139, leading to the formation of a total of 8 hydrogen bonds. These results provide preliminary insights into the potential binding sites of Orthosilicic Acid with the aforementioned two proteins.

## Conclusion

In laboratory culture experiments, Depleted-Repleted silicate conditions were established to investigate changes in silicon accumulation by *Synechococcus* sp. XM24. The Total Biogenic Silica (BSi) content in the Repleted silicate group was significantly higher than in the Depleted silicate group, suggesting cells’ adaptability to varying silicate levels. Silicon content within cells was found to correspond with environmental silicon levels. Notably, the New BSi content in the Depleted silicate group was significantly elevated compared to the Repleted silicate group, indicating an influence of extracellular silicate levels on cellular silicon utilization. Macrotranscriptomic analysis was performed after it was determined that silicate concentration would have a significant effect on the cellular silicon accumulation process. The sequence is screened for 2 genes that appear to *SITs*, which are translated into proteins to predict protein structure and function. The proteins expressed by both sequences may be related to cell transport in Biological processes and cell membrane composition in Cellular components. These results indicate that these 2 genes are most likely related to the silicic acid transport of *Synechococcus* sp. XM24. Molecular docking results identified probable binding sites of the two proteins with Orthosilicic Acid.

## Supporting information

Supplemental Table S1 and Figue S1&S2

## Data availability statement

The data underlying this article will be shared on reasonable request to the corresponding author.

## Author Contributions

Conceptualization: Yabo Han; Methodology: Yabo Han; Writing-Original Draft: Jun Sun and Yabo Han. All authors reviewed the whole work and approved the final version of the manuscript.

## Acknowledgements

This research was financially supported by the National Nature Science Foundation of China grants (41876134), and the State Key Laboratory of Biogeology and Environmental Geology, China University of Geosciences (No.GKZ22Y656).

## Reference

Amo, Y. D. and Brzezinski, M. A. (1999). THE CHEMICAL FORM OF DISSOLVED SI TAKEN UP BY MARINE DIATOMS. J. Phycol. 6,1162–1170. doi: 10.1046/j.1529-8817.1999.3561162.x

Bhattacharyya, P. and Volcani, B. E. (1980). Sodium-dependent silicate transport in the apochlorotic marine diatom Nitz schia alba. Proceedings of the National Academy of Sciences. 11,6386–6390. doi: 10.1073/pnas.77.11.6386

Bäuerlein, E. (2000). Silicic acid transport and its control during cell wall silicification in diatoms. Wiley-VCH Verlag GmbH & Co. KGaA

Hildebrand, M., Volcani, B. E., Gassmann, W. and Schroeder, J. I. (1997). A gene family of silicon transporters. Nature. 6618,688–9. doi: 10.1038/385688b0

Lupas, A., Dyke, M. V. and Stock, J. (1991). Predicting Coiled Coils from Protein Sequences. Science. 5009,1162–1164. doi: 10.1126/science.252.5009.1162

Hildebrand, M., Dahlin, K. and Volcani, B. E. (1998). Characterization of a silicon transporter gene family in Cylindrotheca fusiformis sequences, expression analysis, and identification of homologs in other diatoms. Molecular and General Genetics MGG. 480–486. doi: 10.1007/s004380050920

Armbrust, E. V., Berges, J. A., Bowler, C., Green, B. R., Martinez, D., Putnam, N. H., et al. (2004). The Genome of the Diatom Thalassiosira Pseudonana: Ecology, Evolution, and Metabolism. Science. 5693,79–86. doi: 10.1126/science.1101156

Thamatrakoln, K., Alverson, A. J. and Hildebrand, M. (2006). COMPARATIVE SEQUENCE ANALYSIS OF DIATOM SILICON TRANSPORTERS: TOWARD A MECHANISTIC MODEL OF SILICON TRANSPORT. J. Phycol. 4,822–834. doi: 10.1111/j.1529-8817.2006.00233.x

Alverson, A. J. (2007). Strong purifying selection in the silicon transporters of marine and freshwater diatoms. Limnol. Oceanogr. 4,1420–1429. doi: 10.4319/lo.2007.52.4.1420

Dilkes, B. P., Sapriel, G., Quinet, M., Heijde, M., Jourdren, L., Tanty, V., et al. (2009). Genome-Wide Transcriptome Analyses of Silicon Metabolism in Phaeodactylum tricornutum Reveal the Multilevel Regulation of Silicic Acid Transporters. PLoS ONE. 10,doi: 10.1371/journal.pone.0007458

Curnow, P., Senior, L., Knight, M. J., Thamatrakoln, K., Hildebrand, M. and Booth, P. J. (2012). Expression, Purification, and Reconstitution of a Diatom Silicon Transporter. Biochemistry. 18,3776–3785. doi: 10.1021/bi3000484

Kang, L.-K., Feng, C.-C., Chang, J. and Gong, G.-C. (2015). Diversity and expression of diatom silicon transporter genes during a flood event in the East China Sea. Mar. Biol. 1511–1522. doi: 10.1007/s00227-015-2687-8

Marchenkov, A. M., Bondar, A. A., Petrova, D. P., Habudaev, K. V., Galachyants, Y. P., Zakharova, Y. R., et al. (2016). Unique configuration of genes of silicon transporter in the freshwater pennate diatom Synedra acus subsp. radians. Doklady Biochemistry and Biophysics. 407–409. doi: 10.1134/s1607672916060089

Marchenkov, A. M., Petrova, D. P., Morozov, A. A., Zakharova, Y. R., Grachev, M. A. and Bondar, A. A. (2018). A family of silicon transporter structural genes in a pennate diatom Synedra ulna subsp. danica (Kütz.) Skabitsch. Plos One. 8,doi: 10.1371/journal.pone.0203161

Mann, D. G. (1999). The species concept in diatoms. Phycologia. 6,437–495. doi: 10.2216/i0031-8884-38-6-437.1

Milligan, A. J. and Morel, F. o. M. M. (2002). A Proton Buffering Role for Silica in Diatoms. Science. 5588,1848–1850. doi: 10.1126/science.1074958

Buick, R. (1992). The Antiquity of Oxygenic Photosynthesis: Evidence from Stromatolites in Sulphate-Deficient Archaean Lakes. Science. 5040,74–77. doi: 10.1126/science.11536492

Brasier, M. D., Green, O. R., Jephcoat, A. P., Kleppe, A. K., Kranendonk, M. J. V., Lindsay, J. F., et al. (2002). Questioning the evidence for Earth’s oldest fossils. Nature. 76–81. doi: 10.1038/416076a

Waterbury, J. B., Watson, S. W., Guillard, R. R. L. and Brand, L. E. (1979). Widespread occurrence of a unicellular, marine, planktonic,cyanobacterium. Nature. 293–294. doi: 10.1038/277293a0

Johnson, P. W. and Sieburth, J. M. (1979). Chroococcoid cyanobacteria in the sea: A ubiquitous and diverse phototrophic biomass1. Limnol. Oceanogr. 5,928–935. doi: 10.4319/lo.1979.24.5.0928

Binder, B. J., Chisholm, S. W., Olson, R. J., Frankel, S. L. and Worden, A. Z. (1996). Dynamics of picophytoplankton,ultraphytoplankton and bacteria in the central equatorial Pacific. Deep Sea Research Part II: Topical Studies in Oceanography. 4-6,907–931. doi: 10.1016/0967-0645(96)00023-9

Struyf, E., Smis, A., Van Damme, S., Meire, P. and Conley, D. J. (2009). The Global Biogeochemical Silicon Cycle. Silicon. 207–213. doi: 10.1007/s12633-010-9035-x

Jun, S. and Yuqiu, W. (2018). Preliminary thoughts on Silicon accumulation in Synechococcus. Acta Ecologica Sinica. 14,5234–5243. doi:

Jun, S., Xiaoqian, L., Jianfang, C. and Shujin, G. (2016). Progress in oceanic biological pump. Haiyang Xuebao. 04,1–21. doi:

Richardson, T. L. and Jackson, G. A. (2007). Small Phytoplankton and Carbon Export from the Surface Ocean. Science. 5813,838–840. doi: 10.1126/science.1133471

Lomas, M. W. and Moran, S. B. (2011). Evidence for aggregation and export of cyanobacteria and nano-eukaryotes from the Sargasso Sea euphotic zone. Biogeosciences. 1,203–216. doi: 10.5194/bgd-7-7173-2010

Wei, Y., Sun, J., Li, L. and Cui, Z. (2022). Synechococcus silicon accumulation in oligotrophic oceans. Limnol. Oceanogr. 3,552–566. doi: 10.1002/lno.12015

Conley, D. J., Frings, P. J., Fontorbe, G., Clymans, W., Stadmark, J., Hendry, K. R., et al. (2017). Biosilicification Drives a Decline of Dissolved Si in the Oceans through Geologic Time. Front. Mar. Sci. doi: 10.3389/fmars.2017.00397

Krauskopf, K. B. (1956). Dissolution and precipitation of silica at low temperatures. Geochim. Cosmochim. Acta. 1-2,1–26. doi: 10.1016/0016-7037(56)90009-6

Iler, R. K. (1979). The chemistry of silica: solubility, polymerization, colloid and surface properties, and biochemistry. New York: Wiley-Interscience

Baines, S. B., Twining, B. S., Brzezinski, M. A., Krause, J. W., Vogt, S., Assael, D., et al. (2012). Significant silicon accumulation by marine picocyanobacteria. Nat. Geosci. 12,886–891. doi: 10.1038/ngeo1641

Tang, T., Kisslinger, K. and Lee, C. (2014). Silicate deposition during decomposition of cyanobacteria may promote export of picophytoplankton to the deep ocean. Nat. Commun. 4143. doi: 10.1038/ncomms5143

Deng, W., Monks, L. and Neuer, S. (2015). Effects of clay minerals on the aggregation and subsequent settling of marine Synechococcus. Limnol. Oceanogr. 3,805–816. doi: 10.1002/lno.10059

Guidi, L., Chaffron, S., Bittner, L., Eveillard, D., Larhlimi, A., Roux, S., et al. (2016). Plankton networks driving carbon export in the oligotrophic ocean. Nature. 465–470. doi: 10.1038/nature16942

Ohnemus, D. C., Rauschenberg, S., Krause, J. W., Brzezinski, M. A., Collier, J. L., Geraci-Yee, S., et al. (2016). Silicon content of individual cells of Synechococcus from the North Atlantic Ocean. Mar. Chem. 16–24. doi: 10.1016/j.marchem.2016.10.003

Brzezinski, M. A., Krause, J. W., Baines, S. B., Collier, J. L., Ohnemus, D. C. and Twining, B. S. (2017). Patterns and regulation of silicon accumulation in Synechococcus spp. J. Phycol. 4,746–761. doi: 10.1111/jpy.12545

Krause, J. W., Brzezinski, M. A., Baines, S. B., Collier, J. L., Twining, B. S. and Ohnemus, D. C. (2017). Picoplankton contribution to biogenic silica stocks and production rates in the Sargasso Sea. Global Biogeochem. Cycles. 5,762–774. doi: 10.1002/2017gb005619

Zheng, Q., Wang, Y., Xie, R., Lang, A. S., Liu, Y., Lu, J., et al. (2018). Dynamics of Heterotrophic Bacterial Assemblages within Synechococcus Cultures. Appl. Environ. Microbiol. 3,e01517–17. doi: 10.1128/aem.01517-17

Leblanc, K. and Hutchins, D. A. (2005). New applications of a biogenic silica deposition fluorophore in the study of oceanic diatoms. Limnol. Oceanogr. Methods. 10,462–476. doi: 10.4319/lom.2005.3.462

Azam, F., Hemmingsen, B. B. and Volcani, B. E. (1974). Role of silicon in diatom metabolism: V. Silicic acid transport and metabolism in the heterotrophic diatom Nitzschia alba. Arch. Microbiol. 103–114. doi: 10.1007/BF00403050

Chen, C., Wu, Y., Li, J., Wang, X., Zeng, Z., Xu, J., et al. (2023). TBtools-II: A “one for all, all for one” bioinformatics platform for biological big-data mining. Molecular Plant. 11,1733–1742. doi: 10.1016/j.molp.2023.09.010

Zou, Q., Shen, W., Le, S., Li, Y. and Hu, F. (2016). SeqKit: A Cross-Platform and Ultrafast Toolkit for FASTA/Q File Manipulation. Plos One. 10,e0163962. doi: 10.1371/journal.pone.0163962

Wang, G., Fang, X., Wu, Z., Liu, Y., Xue, Y., Xiang, Y., et al. (2022). HelixFold An Efficient Implementation of AlphaFold2 using PaddlePaddle. arXiv. doi: 10.48550/arXiv.2207.05477

Morris, G. M., Huey, R., Lindstrom, W., Sanner, M. F., Belew, R. K., Goodsell, D. S., et al. (2009). AutoDock4 and AutoDockTools4: Automated docking with selective receptor flexibility. J. Comput. Chem. 16,2785–2791. doi: 10.1002/jcc.21256

Werner, D. (1977). The Biology of diatoms. Berkeley:Univ. of California Press

