## Supplemental Table S1 and Figue S1&S2 for "The silicon transporters (SITs) gene was found in *Synechococcus* sp. XM24"

*Supplementary Material*

Supplemrntary table S1 Artificial sea water medium.

| Component | Final concentration (M) |
| --- | --- |
| Artificial sea water |  |
| NaCl | 4.20×10^-1^ |
| Na_2_SO_4_ | 2.88×10^-2^ |
| KCl | 9.39×10^-3^ |
| NaHCO_3_ | 2.38×10^-3^ |
| KBr | 8.40×10^-4^ |
| H_3_BO_3_ | 4.85×10^-4^ |
| NaF | 7.15×10^-5^ |
| MgCl_2_·6H_2_O | 5.46×10^-2^ |
| CaCl_2_·2H_2_O | 1.05×10^-2^ |
| SrCl_2_·6H_2_O | 6.38×10^-5^ |
| Essential nutrient |  |
| NaH_2_PO_4_·H_2_O | 1.00×10^-5^ |
| NaNO_3_ | 1.00×10^-4^ |
| Na_2_SiO_3_·9H_2_O | 1.00×10^-4^ |
| Trace elements |  |
| CuSO_4_·5H_2_O | 1.96×10^-8^ |
| Na_2_SeO_3_ | 1.00×10^-8^ |
| FeCl_3_·6H_2_O | 1.00×10^-6^ |
| ZnSO_4_·7H_2_O | 7.97×10^-8^ |
| MnCl_2_·4H_2_O | 1.21×10^-7^ |
| CoCl_2_·6H_2_O | 5.03×10^-8^ |
| Na_2_MoO_4_·2H_2_O | 1.00×10^-7^ |
| EDTA | 1.00×10^-4^ |
| Vitamin |  |
| Vitamin H | 2.25×10^-9^ |
| Vitamin B_12_ | 3.70×10^-10^ |
| Vitamin B_1_ | 2.97×10^-7^ |

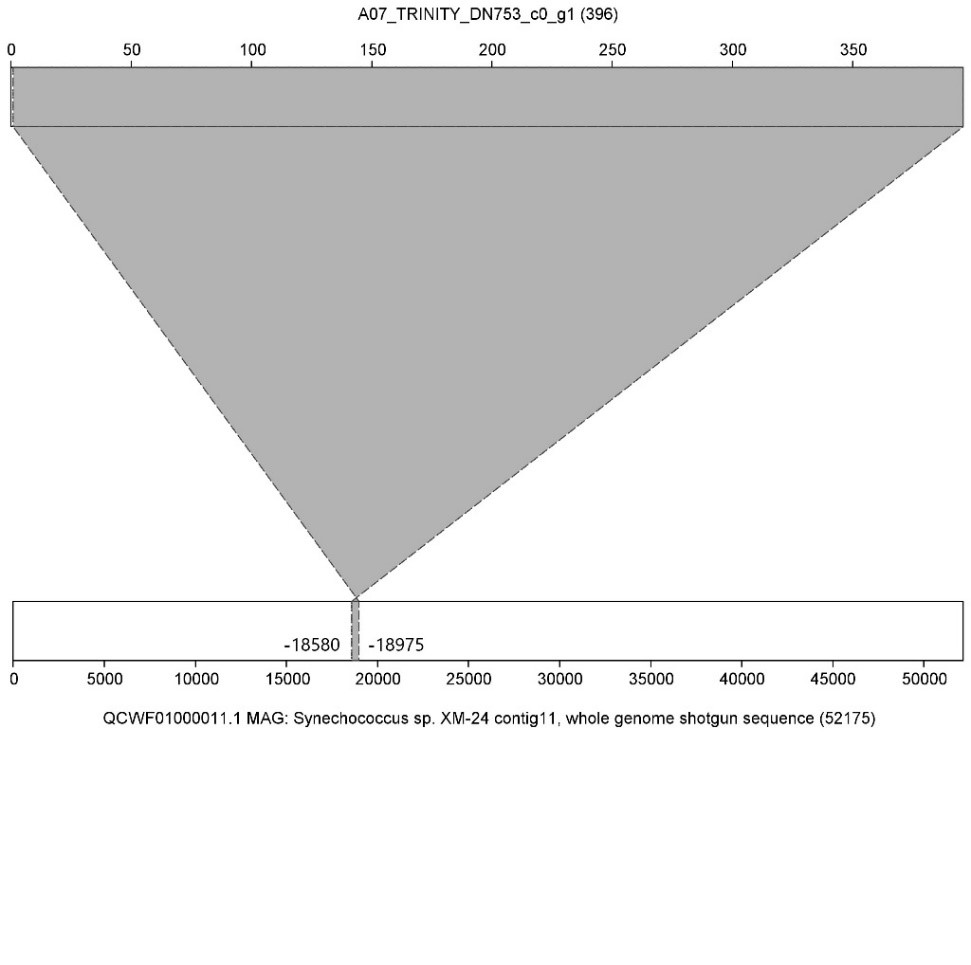

Supplemrntary figure S1 A07_TRINITY_DN753_c0_g1 corresponding to reference genome location.

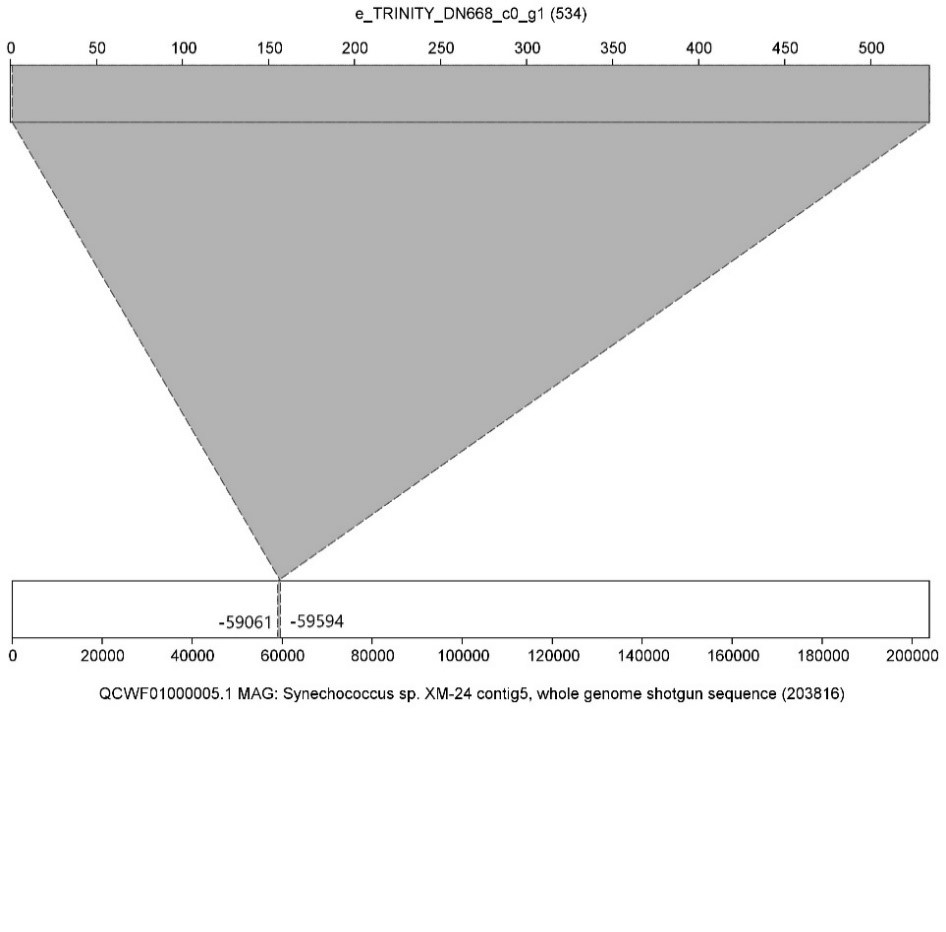

Supplemrntary figure S2 e_TRINITY_DN668_c0_g1 corresponding to reference genome location.
