## Supplementary Materials 2 for "The silicon transporters (SITs) gene was found in *Synechococcus* sp. XM24": Read Me.txt

A07_TRINITY_DN753_c0_g1.fa, e_TRINITY_DN668_c0_g1.fa, e_TRINITY_DN14139_c0_g1.fa—— Gene sequence

OrthosilicicAcid.pdb——The structure file of the Orthosilicic Acid molecule

70percenteTRINITYDN668c0g1transporter——Is the protein structure prediction file after e_TRINITY_DN668_c0_g1 gene is translated into protein, pLDDT is 70%, and the confidence is high

75percenteTRINITY_DN14139c0g1.pdb——Is the protein structure prediction file after e_TRINITY_DN14139_c0_g1 gene is translated into protein, pLDDT is 75%, and the confidence is high

78percentA07TRINITYDN753c0g1transporter.pdb——Is the protein structure prediction file after A07_TRINITY_DN753_c0_g1gene is translated into protein, pLDDT is 78%, and the confidence is high
